# Global lengthening of 3ʹ untranslated regions of mRNAs by alternative cleavage and polyadenylation in cellular senescence

**DOI:** 10.1101/033480

**Authors:** Miao Han, Guoliang Lv, Hongbo Nie, Ting Shen, Yichi Niu, Xueping Li, Meng Chen, Xia Zheng, Wei Li, Chen Ding, Gang Wei, Jun Gu, Xiao-Li Tian, Yufang Zheng, Xinhua Liu, Jinfeng Hu, Wei Tao, Ting Ni

## Abstract

Cellular senescence has been viewed as an irreversible cell cycle arrest that acts to prevent cancer. Recent studies discovered widespread shortening of 3' untranslated regions (3' UTRs) by alternative cleavage and polyadenylation (APA) in cancer cells. However, the role of APA in the process of cellular senescence remains elusive. We thus applied our published PA-seq method to investigate APA regulation in different passages of mouse embryonic fibroblasts (MEFs) and aortic vascular smooth muscle cells (VSMCs) from rats of different ages. We found that genes in senescent cells tended to use distal poly(A) sites (pAs). An independent RNA-seq analysis gave rise to the same conclusion. Interestingly, the level of expression of genes preferred to use distal pAs in senescent MFEs and VSMCs tended to decrease. More importantly, genes that preferred to use distal pAs in senescent MFEs and VSMCs were enriched in common senescence-related pathways such as ubiquitin-mediated proteolysis and cell cycle. Further, the longer 3' UTRs of the genes that tended to use distal pAs introduced more conserved binding sites of senescence-related microRNAs (miRNAs) and RNA binding proteins (RBPs). Noteworthy, the expression level of core factors involved in cleavage and the polyadenylation tended to decrease, while those factors showed opposite trend in cancer cells. In summary, we showed, for the first time, that APA is a hidden layer of post-transcriptional gene expression regulation involved in cellular senescence.

## INTRODUCTION

Cellular senescence was originally described as a process that limits the proliferation of normal human fibroblasts in culture, and causes the loss of telomeres after extensive proliferation in the absence of endogenous telomerase activity (Hayflick and Moorhead 1961; Olovnikov 1996). In fact, in addition to telomere erosion, many stimuli and stress can also cause cellular senescence, including DNA double strand breaks, strong mitogenic signals, oxidative stress, loss of the PTEN tumor suppressor, and ectopic expression of the cyclin-dependent kinase inhibitors (CDKIs) (Rodier and Campisi 2011). Morphologic changes and molecular markers of senescent cells have been identified in the past decades (Munoz-Espin and Serrano 2014). These features include a flatten and enlarged cell morphology, absence of proliferative markers, senescence-associated β-galactosidase (SA-β-gal) activity, expression of tumor suppressors and cell cycle inhibitors, and DNA damage markers (Munoz-Espin and Serrano 2014). Of note, no hallmark of senescence identified thus far is entirely specific to the senescent state, and all senescent cells do not express all possible senescence markers (Munoz-Espin and Serrano 2014).

Cellular senescence is generally viewed as an important mechanism for preventing the growth of cells at risk for neoplastic transformation (Campisi 2001). In recent years, it has become apparent that cellular senescence can also be involved in multiple physiologic and pathologic processes, such as normal embryonic development and tissue damage, suggesting that cellular senescence plays both beneficial and detrimental roles (Munoz-Espin and Serrano 2014). In younger individuals, cellular senescence blocks proliferation of impaired cells, thus preventing transformation into cancer cells and leads to anti-cancer and anti-aging benefits (Munoz-Espin and Serrano 2014). When people get old, however, a growing number of cells enter senescence, which causes decreased tissue function, elevated inflammation, and exhaustion of stem cells, thus becoming pro-aging (Munoz-Espin and Serrano 2014).

A number of studies have shown that dramatic changes in transcriptomes and/or proteomics accompany the dramatic phenotypic changes of senescent cells (Kim YM et al. 2013; Mazin et al. 2013; Waldera-Lupa et al. 2014; Wei et al. 2015). Kim YM et al. profiled cellular senescence phenotypes and mRNA expression patterns during replicative senescence in human diploid fibroblasts (Kim YM et al. 2013). Interestingly, Kim YM et al. reported that the development of the associated senescence phenotypes was supported by the stage-specific gene expression modules (Kim YM et al. 2013). Furthermore, Mazin et al. found widespread splicing changes during human brain development and aging (Mazin et al. 2013). Waldera-Lupa et al. found that 77% of the age-associated proteins were not shown changed message RNA (mRNA) expression level by analyzing an ex vivo model of in situ aged human dermal fibroblasts (Waldera-Lupa et al. 2014). In addition, Wei et al. found decoupling between mRNA level and protein expression modulated by RBPs and miRNAs in prefrontal cortex of elder, suggesting the importance of post-transcriptional regulation via 3' UTR in aging cells (Wei et al. 2015). Further, factors related to different steps of precursor message RNA (pre-mRNA) processing (e.g., splicing, 3' end formation) showed altered level of expression in normal and aged cells (Meshorer and Soreq 2002). These findings raise the possibility that changes in the 3' UTR by APA would contribute to cellular senescence and/or aging.

Cleavage and polyadenylation of nascent RNA is essential for maturation of the vast majority of eukaryotic mRNAs and determines the length of the 3' UTR (Sachs 1990). Such a process requires several cis-acting RNA elements and several dozen core and auxiliary polypeptides (Millevoi and Vagner 2010). The key cis-element that defines cleavage is a 6 nucleotide (nt) motif called the poly(A) signal (PAS), the canonical form of which is AAUAAA (Proudfoot and Brownlee 1976). In addition to the PAS, additional U-rich sequence elements located upstream, as well as U- and GU-rich sequence elements located downstream of the cleavage site can also enhance the efficiency of the 3' end processing reaction (Elkon et al. 2013). The 3' end-processing machinery, contained several sub-complexes, including cleavage and polyadenylation specificity factor (CPSF), cleavage stimulation factor (CstF), cleavage factors I and II (CFI and CFII), as well as additional accessories (Elkon et al. 2013). With the surge in high-throughput sequencing technologies, genomic studies in the past few years have indicated that most eukaryotic mRNA genes have multiple polyadenylation sites (pAs), and thus multiple 3' UTRs caused by APA (Zheng and Tian 2014). Alternative pAs can reside in the 3'-most exon or upstream regions, leading to multiple mRNA isoforms that contain different coding sequences, 3' UTRs, or both (Elkon et al. 2013; Zheng and Tian 2014). Importantly, both miRNAs and RBPs have been reported to control translational efficiency, degradation and subcellular localization of mRNA via targeting to 3' UTRs (Zheng and Tian 2014).

Furthermore, dynamic regulation of 3' UTRs by APA has been reported in different tissue types, development and cellular differentiation, cell proliferation, cell reprogramming, and cancer cell transformation (Zhang et al. 2005; Ni et al. 2013; Wang et al. 2008; Ji et al. 2009; Sandberg et al. 2008; Ji and Tian 2009; Mayr and Bartel 2009; Singh et al. 2009; Xia et al. 2014). Sanberg et al. first reported the link between APA and cell proliferation, indicating fast-growing cells favor proximal pA sites (Sandberg et al. 2008). Consistent with the association between cell proliferation and APA, cancer cells also prefer expression of mRNA isoforms with shorter 3' UTRs compared with normal cells, coupled with increased expression of mRNAs encoding cleavage and polyadenylation (C/P) factors (Mayr and Bartel 2009; Xia et al. 2014). In contrast, progressive 3' UTR lengthening, coupled with weakening of mRNA polyadenylation activity, has been observed during mouse embryonic development and cell differentiation (Ji and Tian 2009). Of note, the regulation of APA is dictated by a combination of several features, including *trans*-acting factors (e.g. C/P factors, tissue-specific RBPs), *cis*-acting elements and epigenomic marks (Di Giammartino et al. 2011).

However, APA regulation in cellular senescence, which regarded as a tumor suppression mechanism, remains unclear. Therefore, we applied our published PA-seq approach, followed by bioinformatic analyses and experimental validations to investigate APA regulation in two different cellular senescence models, including different passages of mouse embryonic fibroblasts (MEFs) and aortic vascular smooth muscle cells (VSMCs) of rats at different ages (Ni et al. 2013). Results showed that many genes tend to use distal pA sites coupled with decreased levels of expression, and also prefer to enrich in shared senescence-related pathways. Interestingly, we found that genes prefer to use distal pAs in cellular senescence surprisingly favor proximal pAs in cancer cells (Xia et al. 2014). We further showed that global 3' UTR lengthening in senescent cells might be explained by decreased expression of cleavage and polyadenylation-related factors. These above results provide the first evidence suggesting that APA is an ignored layer of post-transcriptional gene regulation during cellular senescence.

## Results

### Establishment of a replicative senescence model for MEFs

To uncover the role of APA in cellular senescence, we isolated and continuously sub-cultured MEFs to serve as a typical time-saving replicative senescence model (Busuttil et al. 2003; Parrinello et al. 2003; Di Micco et al. 2008; Manning and Kumar 2010). Population doubling curve showed that MEF cells underwent decreased growth rate (Figure 1A), a traditional marker for replicative senescence. Cells with population doubling (PD) times of 6, 8, 10, and 11 were further analyzed for additional senescence marks. Mki67, a marker for cell proliferation and a robust assessment of cellular senescence (Correia-Melo et al. 2013), showed a decreased level of expression, as determined by quantitative reverse transcription polymerase chain reaction (qRT-PCR; Figure 1B). Flow cytometry revealed a reduced percentage of S phase cells from PD6 to PD11 (Figure 1C). P16, a well-known cyclin-dependent kinase inhibitor that is involved in multiple types of cellular senescence, exhibited elevated expression at both the RNA (Figure 1D) and protein levels (Figure 1E). Senescence-associated β-galactosidase activity (SA-β-gal) also increased from PD6 to PD11 (Figure 1F). All of the above evidence supports the notion that MEF cells from PD6 to PD11 underwent replicative senescence and can be further applied to investigate the potential regulation of APA.

**Figure 1.**
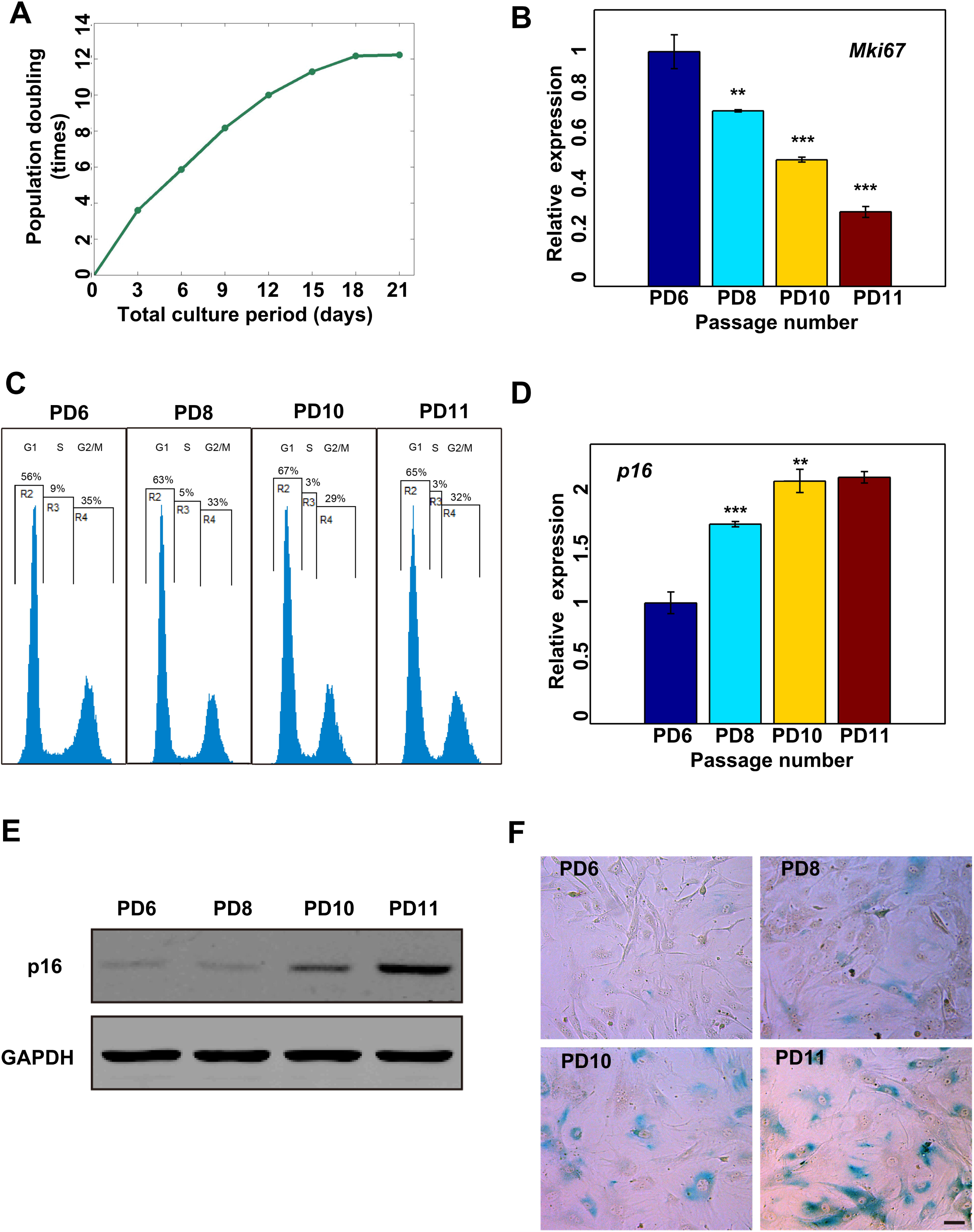
Establishment of a replicative senescence model for MEFs. (A) MEFs were continuously sub-cultured and the population doubling number (passaged number of cells obtained by sequential population doubling) was recorded. (B) Level of expression of *Mki67* in PD6, PD8, PD10, and PD11 MEF passages by qRT-PCR. (C) MEFs form PD6, PD8, PD10, and PD11 passages were subjected to fluorescence-activated cell sorting (FACS) analysis. (D) Level of expression of p16 (encoded by *Cdkn2a*) in PD6, PD8, PD10, and PD11 MEF passages was detected by RT-qPCR. (E) Detection of p16 protein level in PD6, PD8, PD10, and PD11 MEF passages by Western blot analysis. (F) MEFs from PD6, PD8, PD10, and PD11 passages were subjected to senescence-associated β-galactosidase (SA-β-gal) staining. (***) *p <* 0.001 and (**) *P <* 0.01, two tailed Student's t-test.

### Prevalent APA in protein-coding and non-coding transcripts in MEFs

To explore the potential role of APA in replicative senescence, we performed our published PA-seq protocol (Ni et al. 2013) to map the 3' end of mRNAs in 4 different passages of MEF cells (PD6, PD8, PD10, and PD11), which showed multiple senescence markers in later population doubling. To evaluate the influence of the cell cycle phase, which was also associated with cellular senescence, on APA regulation, we investigated the APA of cells in the G0 phase through serum starvation of PD6 cells (the sample is referred to as G0 hereafter). We obtained ~54 million strand-specific paired-end 101-mer reads for the PA-seq, of which ~63% (about ~34 millions) can be uniquely mapped to the mouse reference genome (mm9, NCBI build 37, Supplemental Table S1). The PA-seq data from 4 samples (PD6, PD8, PD10, and PD11), and the G0 stage were combined so that a unified peak-calling scheme could be applied (see Materials and Methods for details). In order to focus the analysis on robustly expressed APA events, we kept pA clusters that accounted for 10% of all the tags within respective genes in any sample and also low-expression pA clusters containing 5% of all the tags within respective genes in the majority of samples (80%). We got 18,175 and 464 high-confident pA clusters in coding genes and non-coding RNAs, respectively, by the above criteria (Figure 2A, Supplemental Table S2). Of these 18,639 pA clusters, 65.8% were included in PolyA_DB 2 (Supplemental Figure S1A), which is a database of pAs in several vertebrate species (Lee et al. 2007). We next examined the genomic distribution of identified pA clusters located in protein-coding genes. Of the pA clusters, 42% were overlapped with known polyadenylation sites (known pA), while 41% and 10% clusters were located in the annotated and extended 3' UTR regions, respectively (Figure 2B). We further evaluated the tag abundance in different categories of pA clusters and found that 56% of all the tags covering the known pAs, while 38% and 4% are located in the annotated and extended 3' UTR regions, respectively (Figure 2C). Further analysis showed that the overall tag number distribution in known pAs also tended to be higher compared to 3' UTR or extended 3' UTR regions (Supplemental Figure S1B). In addition, the median number of covered tags for pAs located in tandem 3' UTR regions was significantly higher than the exon and intron regions (*P*<0.001 and *P*<0.001, Mann-Whitney U test, respectively; Supplemental Figure S1C). The above results indicate that pAs located in known pA sites, 3' UTR, and extended 3' UTR regions are three major groups in MEF cells.

**Figure 2.**
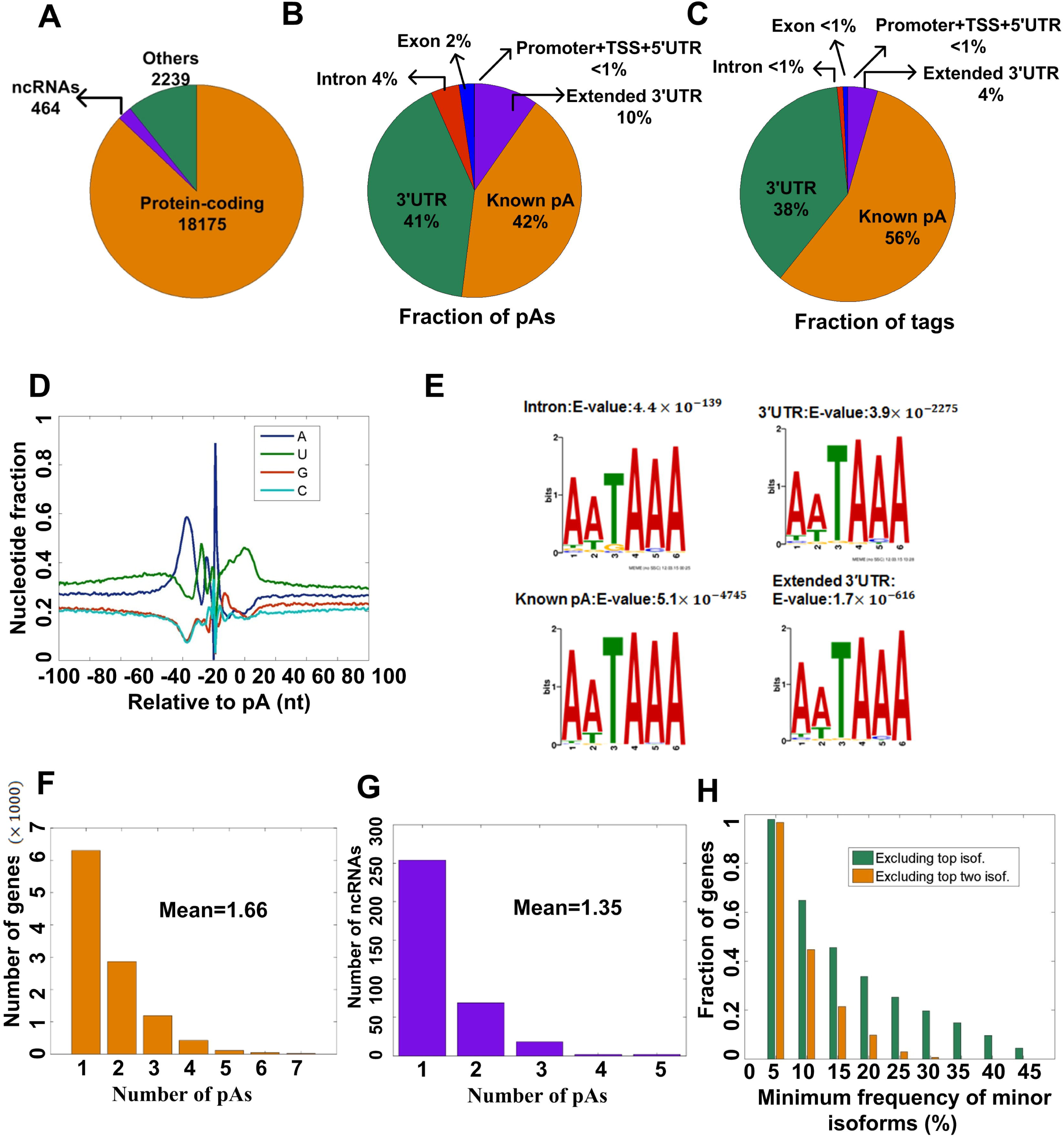
Summary of pAs in MEFs identified by PA-seq. (A) Gene categories of pAs based on RefSeq annotation. (B) The relative locations of the identified pAs in respective genes. (C) The relative locations of the identified pA tags in respective genes. (D) Nucleotide sequence composition around the pAs identified in this study. (E) MEME program identifies the canonical PAS motif, AATAAA, with a very significant E-value from the upstream (-40 bp) of the pAs located in the intron, 3' UTR, known pA, and extended 3' UTR regions. (F) The number of protein-coding genes with a different number of pAs. (G) The number of ncRNAs with a different number of pAs. (H) Fraction of genes with the minimum frequency of minor 3' UTR isoforms exceeding given thresholds (0%, 5%, 15%, 20%, 25%, 30%, 35% 40%, and 45%). Frequencies of minor isoforms were calculated by excluding the top isoform (green bars) and the top two isoforms (orange bars).

The nucleotide composition near the pAs was analyzed to further validate the reliability of those identified cleavage and polyadenylation sites. In agreement with a previous study (Hoque et al. 2013), adenine base (A) had a higher fraction at -40 to -20 nucleotides (nt) before the pA sites, while uridine base (U) peaks downstream of the pA sites (Figure 2D). Furthermore, the nucleotide preference downstream the site of cleavage is in the order of A>U>C>G for pAs located in different regions of genes, which also in line with previous study (Figure 2D and Supplemental Figure S1D) (Chen et al. 1995). In addition, all of the pAs identified in this study, except for the pAs located in the exon, shared very similar features: canonical PAS was found upstream of the pAs; there was a relatively high U content and lower C/G content surrounding the pAs and also high frequencies of *cis* elements defined by polya_svm (Supplemental Figure S1E-G) (Cheng et al. 2006).

Alternative polyadenylation sites were further investigated for both protein-coding genes and non-coding RNAs (ncRNAs). Our PA-seq analyses revealed that ~42% of all expressed protein-coding genes have two or more pAs (Figure 2F, Supplemental Figure S2A-B) and each gene, on average, had 1.66 APA isoforms, which is comparable with another study based on PAS-seq (Shepard et al. 2011). In addition, we identified 464 pAs covering 345 ncRNAs, and 26% of these ncRNAs had two or more pAs and the average number of pAs was 1.35 (Figure 2G, Supplemental Figure SC). We found remaining isoforms other than top expressing ones also showed considerable abundance for genes with multiple pAs (Figure 2H). Therefore, multiple pAs should be considered at the same time to avoid systematic bias that could be introduced if only two pA sites are considered.

Based on the above analyses, the top three categories of pAs (known pA, 3' UTR, and extended 3' UTR) occupied 93% (Figure 2B) of all identified pAs and 98% (Figure 2C) of tags inside all of the identified pA clusters. Therefore, the combined tags in these three pA clusters (number of Tags Per Million reads [TPM]) were used to evaluate the level of expression of each gene with APA. To examine whether or not the PA-seq results reflect the relative level of gene expression, the same RNA samples were sequenced by strand-specific RNA-seq (Parkhomchuk et al. 2009). In total, we obtained ~104 million paired-end 101-mer reads, and approximately 92% of the reads (~96 millions) were mapped uniquely to the mouse genome (Supplemental Table S3). Specifically, we calculated the gene expression from the RNA-seq data with the fragment per kilobase of exon model per million (FPKM) value (Trapnell et al. 2012). We only considered genes expressed in all passages (FPKM ≥ 1 and TPM ≥ 1) for further analysis. PA-seq and RNA-seq profiles were considerably correlated (Spearman's rank correlation coefficient rho = 0.81, 0.83, 0.79, and 0.78 for PD6, PD8, PD10, and PD11, respectively; Supplemental Figure 3 and Supplemental Table S4). Therefore, the read count generated by the PA-seq approach can potentially be used to reflect transcript abundance.

**Figure 3.**
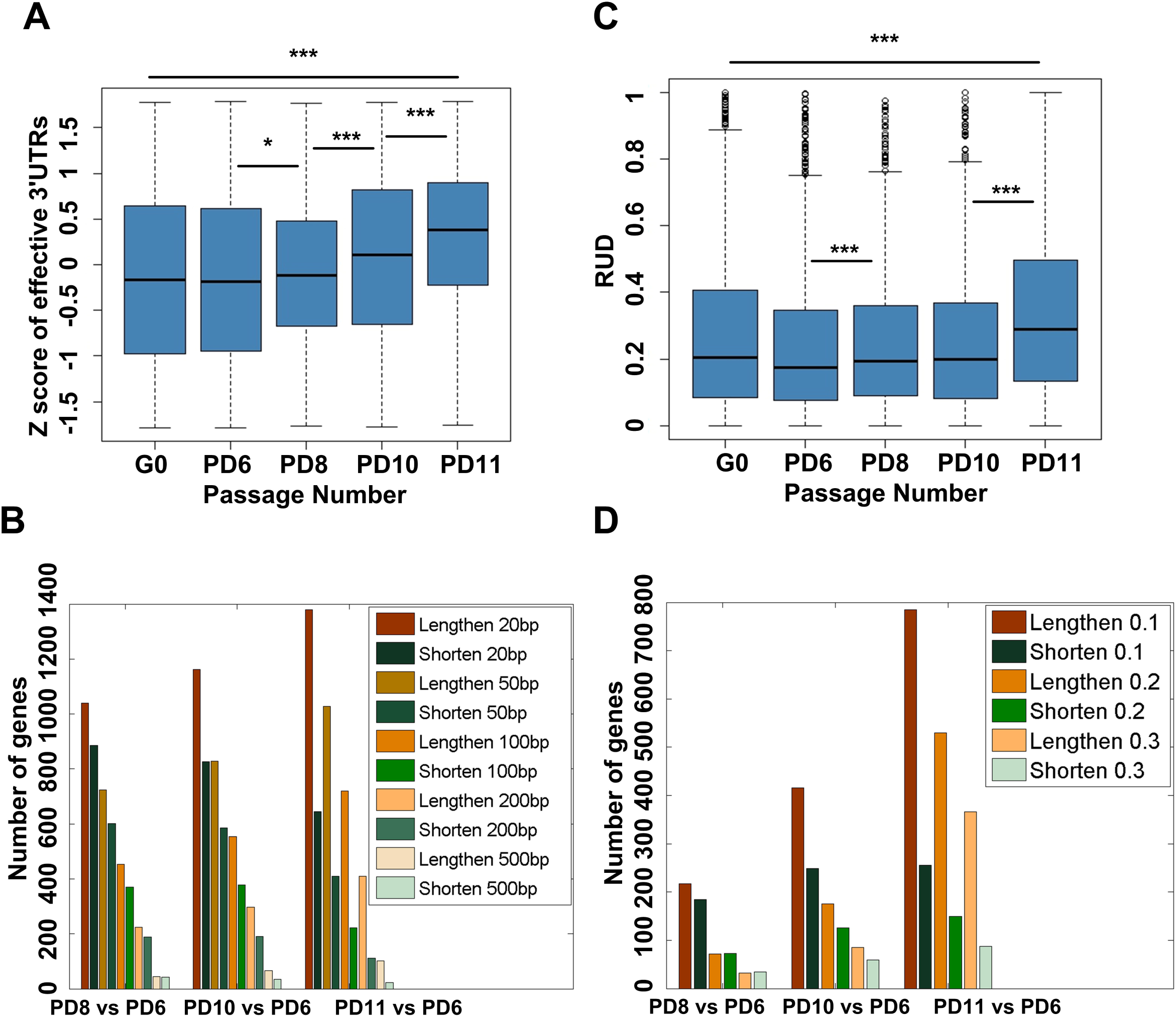
Global lengthening of 3' UTRs for genes with APA regulation during replicative senescence of MEFs. (A) Box plot for Z scores of effective 3' UTRs across G0, PD6, PD8, PD10, and PD11 of MEFs. (B) Number of genes with lengthened effective 3' UTRs and number of genes with shortened effective 3' UTRs by comparing PD11, PD10, and PD8 to PD6 given different thresholds based on PA-seq data. (C) Box plot for relative expression of mRNA isoforms using distal pA sites (RUD scores) across G0, PD6, PD8, PD10, and PD11 of MEFs. (D) Number of genes with higher RUD scores and number of genes with lower RUD scores by comparing PD11, PD10, and PD8 to PD6 given different cutoffs based on RNA-seq data. (***) *P* < 0.001 and (*) *P* < 0.05, two-tailed Wilcoxon signed rank test.

### Global lengthening of 3' UTRs for genes with APA during replicative senescence of MEFs

Because our PA-seq data contained relatively reliable pAs, as well as the steady state level of expression, a unique opportunity was provided to interrogate global changes of 3' UTR length of APA genes among different passages of MEF cells. Based on our previous study (Ni et al. 2013), the effective 3' UTR length (effective_UTR hereafter) was used to estimate the relative use of pAs inside or downstream of 3 ' UTR regions for each gene based on a consolidated gene model by employing the locations and the tag counts of the p A clusters identified in the 3 ' UTR of each transcribed locus (Ni et al. 2013). We are excited to find that effective 3' UTR showed a global progressive lengthening trend during the population doubling process of MEF cells (Figure 3A, Supplemental Table S4). To further evaluate the changes in 3' UTR length at the individual gene level, we compared the effective 3' UTR length in later passages (PD8, PD10, and PD11) with an earlier passage (PD6) using different cut-offs (20bp, 50bp, 100bp, 200bp, and 500bp; Figure 3B). There were always an increased number of genes with a longer effective 3' UTR region than a shortened effective 3' UTR region when compared to PD6 at different thresholds (Figure 3B). Of note, the number of genes with a lengthened effective 3 ' UTR region was gradually increased from PD8 to PD11 when compared to PD6, while the number of genes with a shortened effective 3' UTR region was continuously decreased from PD8 to PD11 when compared to PD6 (Figure 3B). These results further validated the global lengthening trend during the process of replicative senescence, as shown in Figure 3A.

In agreement with the PA-seq data, we also observed the same trend of global 3' UTR lengthening during replicative senescence in MEFs using measurement of the RUD index based on RNA-seq data by following the analysis pipeline of Zhe et al. (Ji et al. 2011) (Figure 3C, Supplemental Table S4). The change in RUD index between different passages was applied to evaluate the preference of a longer or shorter 3' UTR region for specific genes. In agreement with PA-seq results, more genes have a longer 3' UTR region in later passages (PD8, PD10, and PD11) than in an earlier passage (PD6; Figure 3D). In addition, the number of genes with a higher RUD index (labeled “lengthen” in Figure 3D) was continuously elevated from PD8–11 when compared to PD6, while the number of genes with a decreased RUD index (labeled “shorten” in Figure 3D) was gradually reduced from PD8–11 when compared to PD6 (Figure 3D). This independent RUD analysis confirmed the global lengthening of 3' UTRs during replicative senescence in MEF cells.

To determine whether or not the cell cycle affects the effective 3' UTR length, MEF cells at population doubling time 6 (PD6) were treated with serum starvation to force them entry into the G0 phase, a reversible cell cycle arrest state. We found the global 3' UTR pattern of the G0 sample was closer to PD6 than PD11, a more senescent status, by both PA-seq (Figure 3A) and RNA-seq (Figure 3C) data. This result implies that global lengthening of 3' UTR is more specific to cellular senescence (or irreversible cell cycle arrest) than to reversible cell cycle arrest (G0 phase).

To validate that genes did undergo 3' UTR lengthening during cellular senescence, we randomly selected 10 genes for qRT-PCR validation (Figure S4-S5; see details about primers in Supplemental Table S5). Two genes that had a shortened 3' UTR region in PD11 compared to PD6 were included as internal controls (Supplemental Figure S5). All 10 genes showed an elevated trend of using a longer 3' UTR region (Supplemental Figure S5), with 7 genes reaching a statistically significant level (*P*≤ 0.05, two-tailed Student's t-test). As a confirmation that the qRT-PCR approach reflected the trend towards a change in 3' UTR length, the two control genes showed a shortened 3' UTR usage in PD11 compared to PD6, with statistical significance (P≤ 0.05, two-tailed Student's t-test; Supplemental Figure S5). The experimental validation supports the observation that senescent cells underwent global 3' UTR lengthening.

### Longer 3' UTRs tend to have a decreased abundance of mRNA during cellular senescence

We next examined the consequence of 3' UTR lengthening during MEF cellular senescence. A longer 3' UTR region is thought to provide more opportunities for regulation by miRNAs and/or RBPs, which affects the abundance of mRNA and/or translation at the post-transcriptional level (Zheng and Tian 2014). In line with this hypothesis, a global decrease in gene expression was observed during cellular senescence for genes that have multiple pAs (Figure 4A). Internal control genes with a single pA site did not show such trend (Figure 4B), which supporting the notion that global down-regulation of the level of expression is specific for APA genes. A change in the expression at the individual gene level was also analyzed. To simplify, we compared PD11 MEF cells with PD6 MEF cells for 3' UTR usage and the steady state level of expression. The genes that showed a significantly lengthened 3' UTR outnumbered genes containing a shortened 3' UTR four to one (FDR ≤ 0.05, linear trend test; Figure 4C, Figure S6A, and Supplemental Table S7). Within 3' UTR lengthened genes, a greater number of genes showed a significant down-regulation than genes with an elevated level of expression (*P* < 5.6 × 10^−12^, Binomial test; Figure 4C, right half). No such difference was observed for genes with a shorter 3' UTR (Figure 4C, left half). Together, decreased expression at the individual gene level supports the observation of global down-regulation of APA genes during cellular senescence.

**Figure 4.**
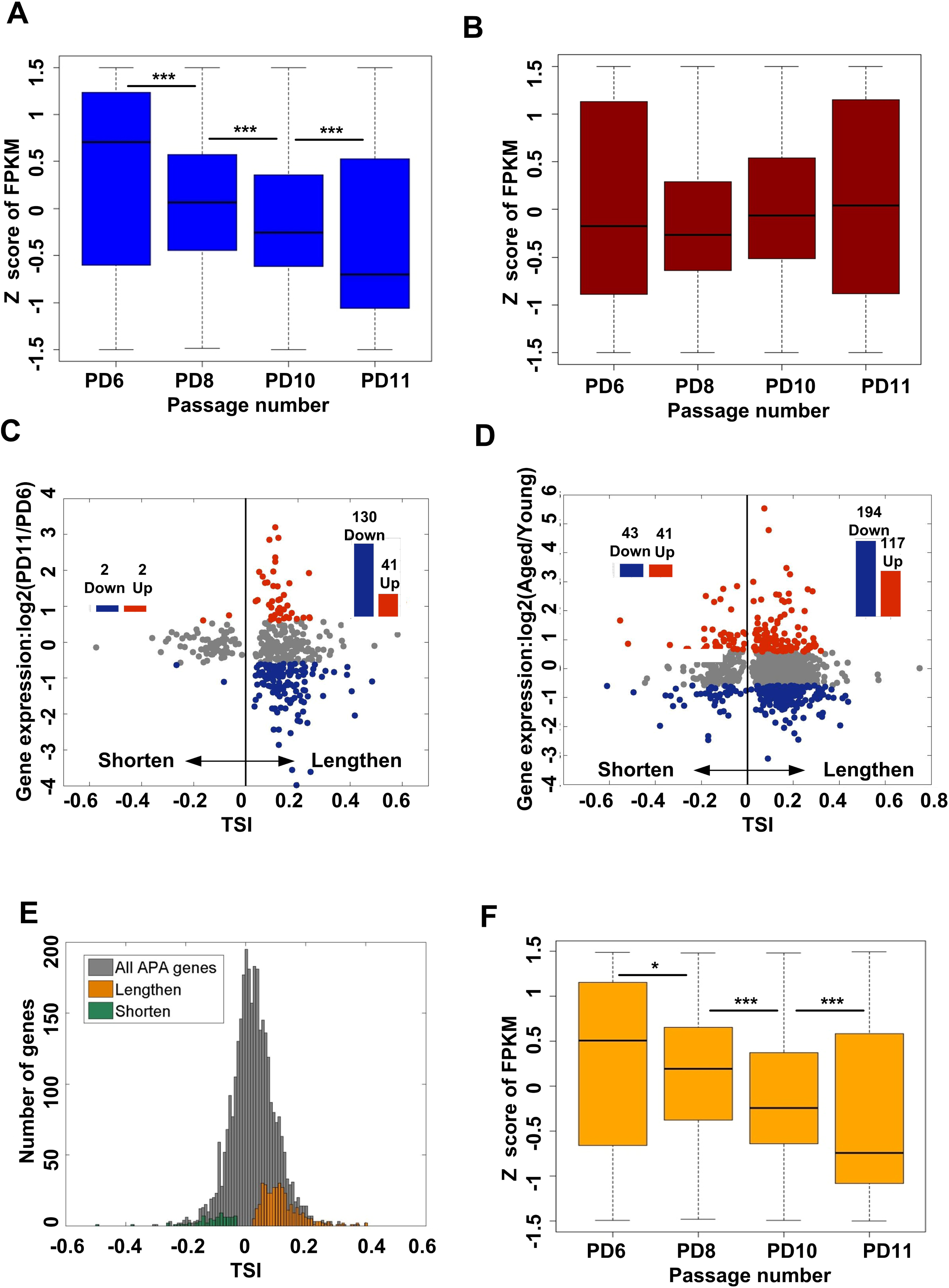
Genes preferred to use distal pAs in senescent cells tended to have decreased mRNA abundance. (A) Box plot of FPKM values across PD6, PD8, PD10, and PD11 for genes with multiple pAs. (B) Box plot of FPKM values across PD6, PD8, PD10, and PD11 for genes with single pA site. (C) For genes with significantly longer or shorter 3' UTRs in senescent MEFs (PD11), the fold-change (FC) expression between senescent MEFs and younger MEFs (PD6) was plotted against the tandem UTR isoform switch index (TSI) values (see Materials and Methods for details). Red and blue represented up- and down-regulation (fold-change ≥1.5 and FPKM ≥1 in both samples) in senescent cells, respectively. The genes with significantly longer or shorter 3' UTRs in senescent MEFs were identified by a linear trend test with the Benjamini-Hochberg (BH) false-discovery rate at 5%. (D) For genes with significantly longer or shorter 3' UTRs in VSMCs derived from old rats (2 years old), the fold-change expression between VSMCs of old and young rats were plotted against the TSI values. Red and blue represented up- and down-regulation (fold-change > 1.5 and TPM > 1 in both samples) in senescent cells, respectively. (E) Distribution of TSI values for all APA regulation genes. Orange and green bars show the distribution of TSI values for genes using lengthened and shortened 3' UTRs during replicative senescence of MEFs, respectively. (F) Box plot of FPKM values across PD6, PD8, PD10, and PD11 for genes with progressively lengthened 3' UTRs during replicative senescence of MEFs. (***) *p* < 0.001 and (*) *P* < 0.05, two-tailed Wilcoxon signed rank test.

To address whether or not APA regulates gene expression in another aging system, we constructed PA-seq libraries for total RNA from vascular smooth muscle cells (VSMCs) derived from young and aged rats. PA-seq libraries were sequenced and analyzed using the same strategy (see read mapping information in Supplemental Table S6). Unexpectedly, a very close ratio of longer versus shorter genes regarding 3' UTR length between rat VSMCs and mouse MEF cells was observed (Figure S6A, Supplemental Table S7). Moreover, a significant overlap of genes showed longer 3 ' UTR usage between mouse MEFs and rat VSMCs (*P* < 1 × 10^−3^, Fisher's exact test; Figure S7-S8). Further, the correlation that more genes have down-regulated expression given lengthened 3' UTR also occur in rat VSMCs (*P* < 1.5 × 10^−5^, Binomial test; Figure 4D). The above results support the notion that APA-medicated 3' UTR lengthening may play a role in gene expression in multiple cellular senescence systems.

To further determine whether or not there is a continuous lengthening of 3' UTR at the individual gene level and the effect on gene expression, a linear trend test was further used to identify genes with progressive 3 ' UTR length changes in MEFs (Fu et al. 2011; Li et al. 2012). Three hundred seventy-five genes showed a continuous lengthening of 3' UTR from PD6–11, while only 73 genes had a progressively shortened 3' UTR (Figure 4E; see details in Supplemental Table S7). Interestingly, a similar global decrease in expression of these 375 genes correlated with the continuously lengthened 3' UTR (Figure 4F, Figure S6B). Noteworthy, the 73 genes contained a continuously shortened 3' UTR during cellular senescence did not display a trend towards expression of down-regulation (Figure S6C-D). Together, the above results indicated that global 3' UTR lengthening may contribute to a decreased mRNA steady state level of expression during cellular senescence.

### Functional enrichment analysis of genes with longer 3' UTRs during cellular senescence

To address whether or not 3' UTR lengthening is related to regulation of the process of cellular senescence, genes with a longer 3' UTR were analyzed for functional enrichment. We first selected genes that showed lengthening in PD11 compared to PD6 since these two passages of MEFs displayed dramatic phenotypic changes (Figure 1F). The Database for Annotation, Visualization, and Integrated Discovery (DAVID) was applied for pathway enrichment analysis because this tool is widely used (Huang et al. 2009). For the 322 genes that have a significantly longer 3' UTR in PD11 compared with PD6 of MEFs, the top-enriched pathways have connections to cellular senescence (Figure 5A, Supplemental Table S8). Genes that progressively tend to use distal pAs during replicative senescence of MEFs are also enriched in senescent-related pathways (Figure S9, Supplemental Table S8). Intriguingly, most of the top-enriched pathways in rat VSMCs are also related to cellular senescence (Figure 5B, Supplemental Table S8). Four of the pathways (marked in red for Figure 5A and 5B) are shared between mouse MEFs and rat VSMCs, including ubiquitin-mediated proteolysis, the wnt signaling pathway, cell cycle, and regulation of the actin cytoskeleton, all linked to cellular senescence ( Deschenes-Simard et al. 2014; Hofmann et al. 2014; Chandler and Peters 2013; Amberg et al. 2012;) (see Discussions for details). We further examined whether or not these two species (mouse and rat) not only shared the same pathways, but the same genes or the genes themselves underwent APA regulation in a conservative fashion. Of note, six genes involved in ubiquitin-mediated proteolysis showed a longer 3' UTR in both mouse and rat senescent cells (Figure 5C). Seven, 2, and 5 genes involved in the regulation of the actin cytoskeleton, cell cycle, and wnt signaling pathway, respectively (Figure 5D-F), had the same trend towards 3' UTR lengthening in both mouse and rat senescent cells. The above functional enrichment results supported the idea that an intrinsic link exists between APA genes and cellular senescence.

**Figure 5.**
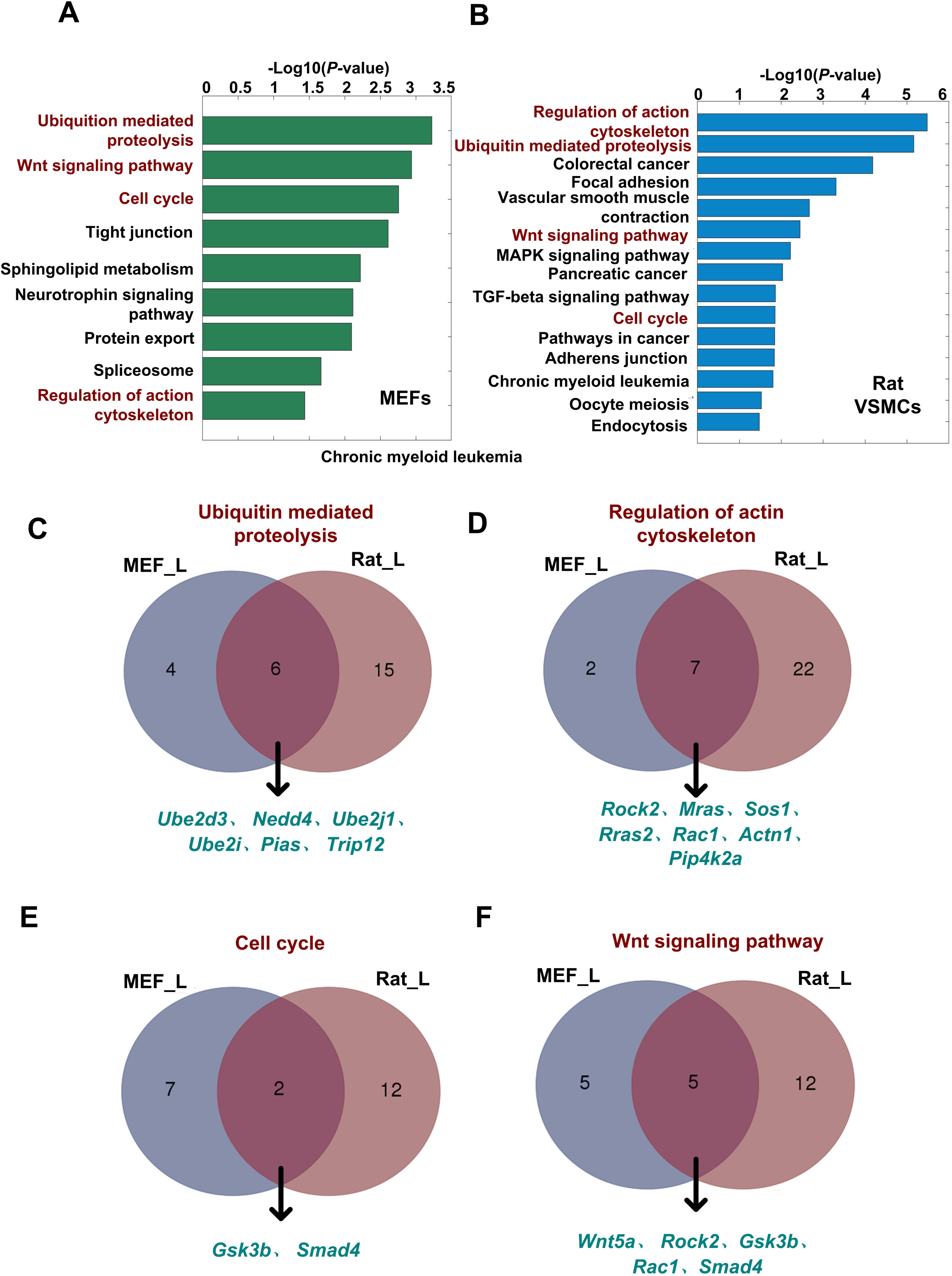
Genes preferred to use distal pAs both in senescent MEFs and rat VSMCs are enriched in shared senescence-related pathways. (A-B) Significantly enriched (*P* <0.01, Fishers' exact test) pathways of the genes tended to use distal pAs in senescent MEFs and VSMCs derived from old rats. The shared senescence-related pathways are marked in red. (C-F) Venn diagram comparison of genes, which preferred to use distal pAs between senescent MEFs and VSMCs of old rat, involved in ubiquitin-mediated proteolysis, regulation of the actin cytoskeleton, cell cycle, and the wnt signaling pathway.

### Longer 3' UTRs introduced more conserved miRNA binding sites and RBP recognition sequences

To explain why a longer 3' UTR decreases the level of mRNA expression during cellular senescence, we analyzed the number and density of recognition sites by miRNA and RBP in the alternative 3' UTR regions between promoter proximal and distal pA sites. In line with previous studies, we found that the number of conserved miRNA binding sites in long 3' UTR (determined by distal pA site [3' UTR_L hereafter]) and alternative 3' UTR (between proximal and distal pA site [3' UTR_A hereafter]) were significantly greater than in short 3' UTR (determined by proximal pA site [3' UTR_S hereafter]) region (*P<* 2.2 × 10^−16^ and *P<* 5.6 × 10^−7^, two-sided Wilcoxon signed rank test, respectively; Figure 6A) (Sandberg et al. 2008). Notably, 3' UTR_L and 3' UTR_A were also significantly longer than 3' UTR_S (P< 2.2 × 10^−16^ and *P<* 2.2 × 10^−16^, two-sided Wilcoxon signed rank test, respectively; Figure 6B). Furthermore, the conserved miRNA binding density in 3' UTR_S was significantly greater than 3' UTR_L and 3' UTR_A (P< 2.2 × 10^−16^ and *P<* 2.4 × 10^−16^, two-sided Wilcoxon signed rank test, respectively; Figure 6C). However, some conserved miRNA binding sites were exclusively located in alternative 3' UTR (Figure 6D). Interestingly, there was a higher density of binding sites in 3' UTR_L than 3' UTR_S for mir-20a-5p, which had been confirmed to be involved in cellular senescence of MEFs (Poliseno et al. 2008) (*P<* 0.03, two-sided Wilcoxon signed rank test; Figure 6E). Similarly, miR-290a-5p, which acts as a physiologic effector of senescence in MEFs (Pitto et al. 2009), also had increased conserved binding density in 3' UTR_L and 3' UTR_A than in 3' UTR_S (*P*< 2.5 × 10^−3^ and *P<* 9.4 × 10^−3^, two-sided Wilcoxon signed rank test, respectively; Figure 6F). These above results indicate more potential miRNA binding sites can be introduced in longer 3' UTR, although the density of binding can be either high or low in the alternative region.

**Figure 6.**
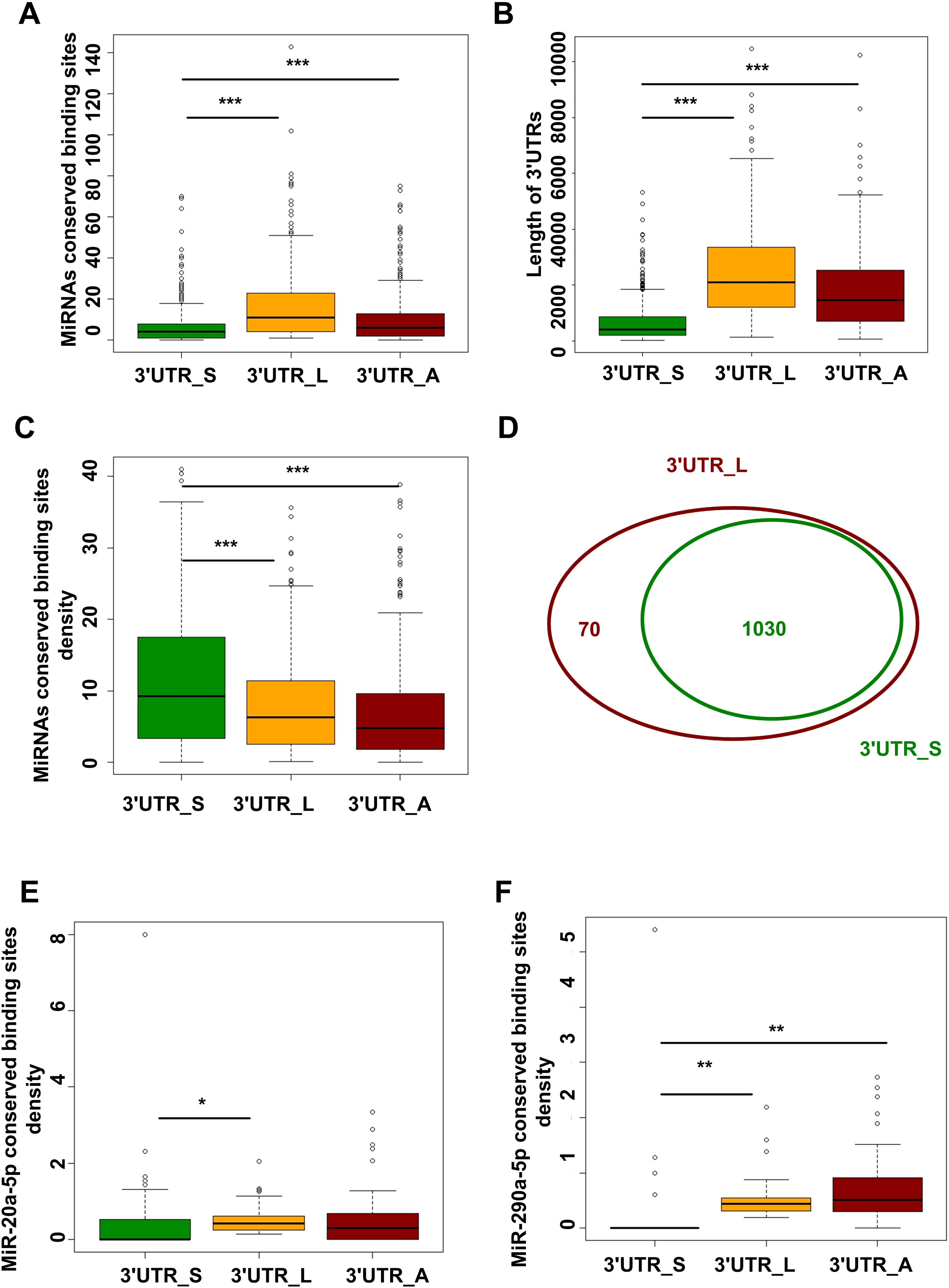
Genes progressively tend to use distal pAs during replicative senescence of MEFs can introduce more conserved miRNA binding sites. (A) Box plot comparison for the number of conserved miRNA binding sites among the shortest 3 ' UTRs (3'UTR_S), the longest 3' UTRs (3'UTR_L), and alternative 3' UTRs (3'UTR_A) in genes that tend to progressively use distal pAs during replicative senescence of MEFs. (B) Box plot for length comparison among 3' UTR_S, 3'UTR_L and 3'UTR_A. (C) Box plot comparison for density of conserved miRNA binding sites among 3' UTR_S, 3'UTR_L, and 3'UTR_A. (D) Venn diagram comparing conserved miRNA binding sites in the 3' UTR_S with 3' UTR_L. (E) Comparing the density of miR-20a-5p potential binding sites in 3' UTR_S with 3' UTR_L and 3' UTR_A. (F) Comparing the density of miR-290a-5p potential binding sites in 3' UTR_S with 3' UTR_L and 3' UTR_A. (***) *P* <0.001, (**) *P* <0.01 and (*) *P*<0.05, two-tailed Wilcoxon signed rank test.

The RNA-Binding Protein DataBase (RBPDB) is a collection of experimentally validated RNA-binding sites both *in vitro* and *in vivo*, or manually curated from the primary literature (Cook et al. 2011). Interestingly, we found that approximately 58% expressed RBPs (307 expressed RBPs with a FPKM > 1 in PD6, PD8, PD10, and PD11), as annotated by this database, was decreased during replicative senescence of MEFs (Figure 7A). RBPmap can predict the binding sites of 94 human/mouse RBPs, 59 among which are also annotated by RBPDB (Paz et al. 2014). Using this tool, we found that 30 expressed RBPs had significantly higher conserved binding densities in both 3' UTR_L and 3' UTR_A than in 3' UTR_S (FDR < 0.05, two-sided Wilcoxon signed rank test; Supplemental Table S9). Specifically, HNRNPA1 (Roy et al. 2014) and IGF2BP2 (Nielsen et al. 2001) can be involved in mRNA translocation, YBX1 (Capowski et al. 2001) can regulate mRNA stability, and NOVA1 (Licatalosi et al. 2008), SNRNP70 (Gunderson et al. 1998), HNRNPF (Decorsiere et al. 2011), HNRNPH2 (Bagga et al. 1998), and HNRNPK (Naganuma et al. 2012) can regulate APA (Figure 7B-J). Taken together, the above data indicated that longer 3' UTRs introduced more sites for RNA binding proteins, which may further affect mRNA stability, localization, and even contribute to alternative polyadenylation.

**Figure 7.**
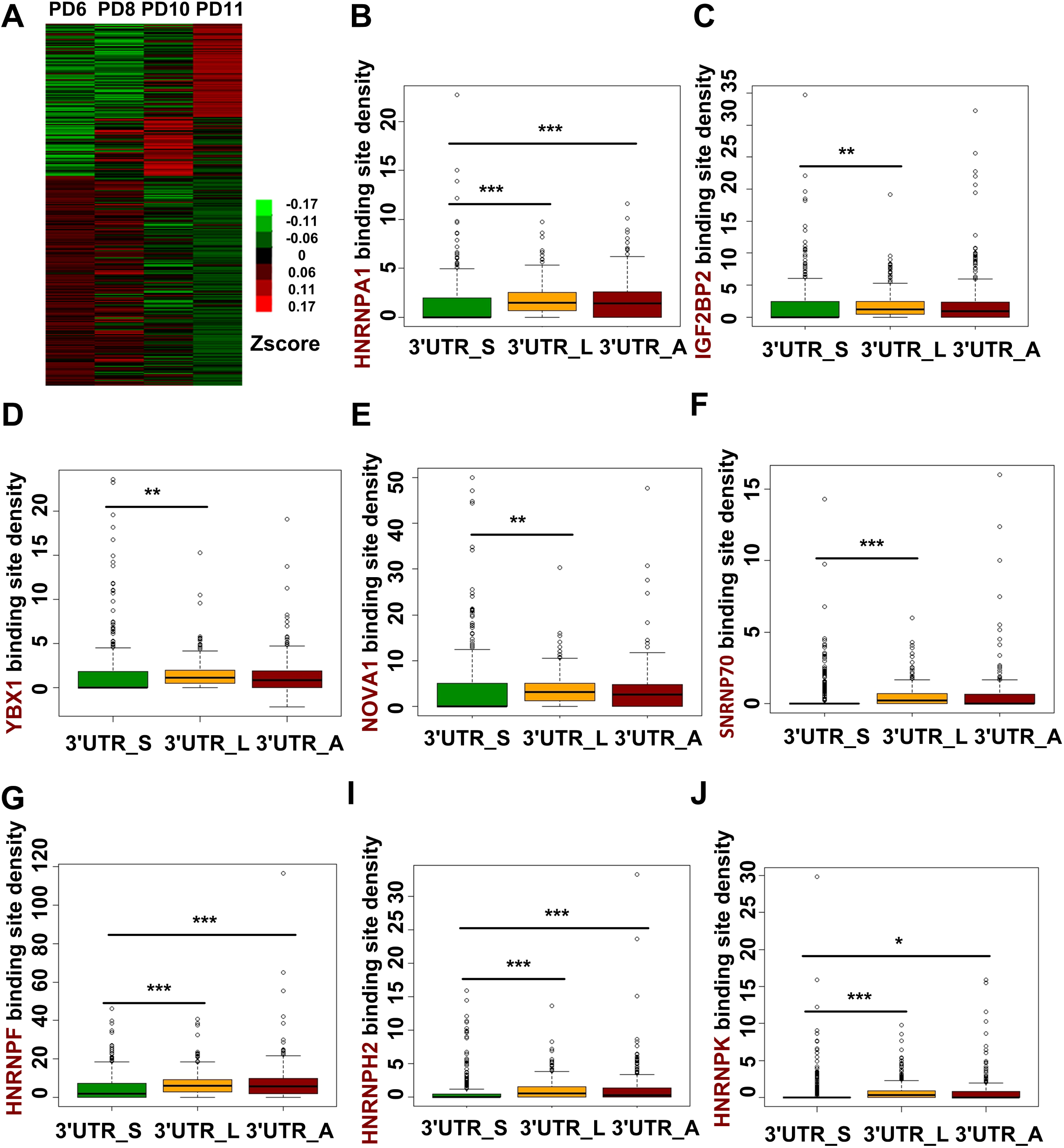
Genes with progressively lengthened 3' UTRs during replicative senescence of MEFs can introduce more conserved RBPs binding sites. **(A)** Heat map of RBPs expression during replicative senescence of MEFs. **(B-J)** Comparing the potential binding site density of HNRNPA1, IGF2BP2, YBX1, NOVA1, SNRNP70, HNRNPF, HNRNPH2, and HNRNPK in the shortest 3' UTRs (3'UTR_S) with the longest 3' UTRs (3'UTR_L) and alternative 3' UTRs (3'UTR_A). (***) *P* <0.001, (**) *P* <0.01 and (*) *P* <0.05, two-tailed Wilcoxon signed rank test.

### Decreased expression of cleavage and polyadenylation-related factors may regulate global 3' UTR lengthening

There are three types of regulators that control alternative polyadenylation, including trans-acting RBPs, cis-acting elements, and chromatin features near pA sites (Di Giammartino et al. 2011). *Cis* elements around proximal or distal pA sites were first analyzed to get insight regarding APA regulation during cellular senescence. We compared the sequence conservation around the regions of the proximal and distal pAs of genes that showed progressive lengthening of 3' UTR (LE_proximal and LE_distal, respectively) to the other three groups of pA sites, including proximal and distal pAs of genes that showed progressive shortening of 3' UTR (SH_proximal and SH_distal, respectively), proximal and distal pAs of genes without significant changes of pAs usage preference (NC_proximal and NC_distal, respectively), and pA regions of genes only contain one pA site (single pA). Interestingly, the polyadenylation signal (PAS) upstream of LE_distal pAs was more conserved than LE_proximal pAs (Figure 8A), and the percentage of ‘AAUAAA’, the strongest PAS, in LE_distal pA sites was the highest among all these seven types of pA sites (Figure 8B). In addition, the downstream sequence conservation scores of proximal pA sites (including LE_proximal, SH_proximal, and NC_proximal) were clearly higher than distal pA sites (including LE_distal, SH_distal, and NC_distal) (Figure 8A). This finding may be attributed to lower frequencies of strong PAS in proximal pAs than distal pAs, therefore more binding sites for RBPs are needed (Figure 8B). Further, among three types of proximal pA sites, both the upstream and downstream sequence conservation score of pAs with a significant usage shift (LE_proximal, SH_proximal) were more conserved than pAs without a significant usage shift (NC_proximal) (Figure 8A). This finding may also be attributed to more binding sites for RBPs needed to help the pAs with a significant usage shift to be used. In agreement with this hypothesis, a global less conservation score of single pAs without APA regulation, except for a higher conservation score in PAS regions, was observed surrounding the pAs (Figure 8A). In addition, both the upstream and downstream sequence conservation score of LE_proximal was higher than SH_proximal (Figure 8A). Those above results raise the possibility that genes showed lengthening of 3' UTR during cellular senescence was regulated by the decreased abundance or activity of RBPs, which can bind with the more conserved sequence around LE_proximal pAs and are necessary for LE_proximal pAs to be used, therefore leaded to increase usage of LE_distal pAs with more conserved and stronger PAS.

**Figure 8.**
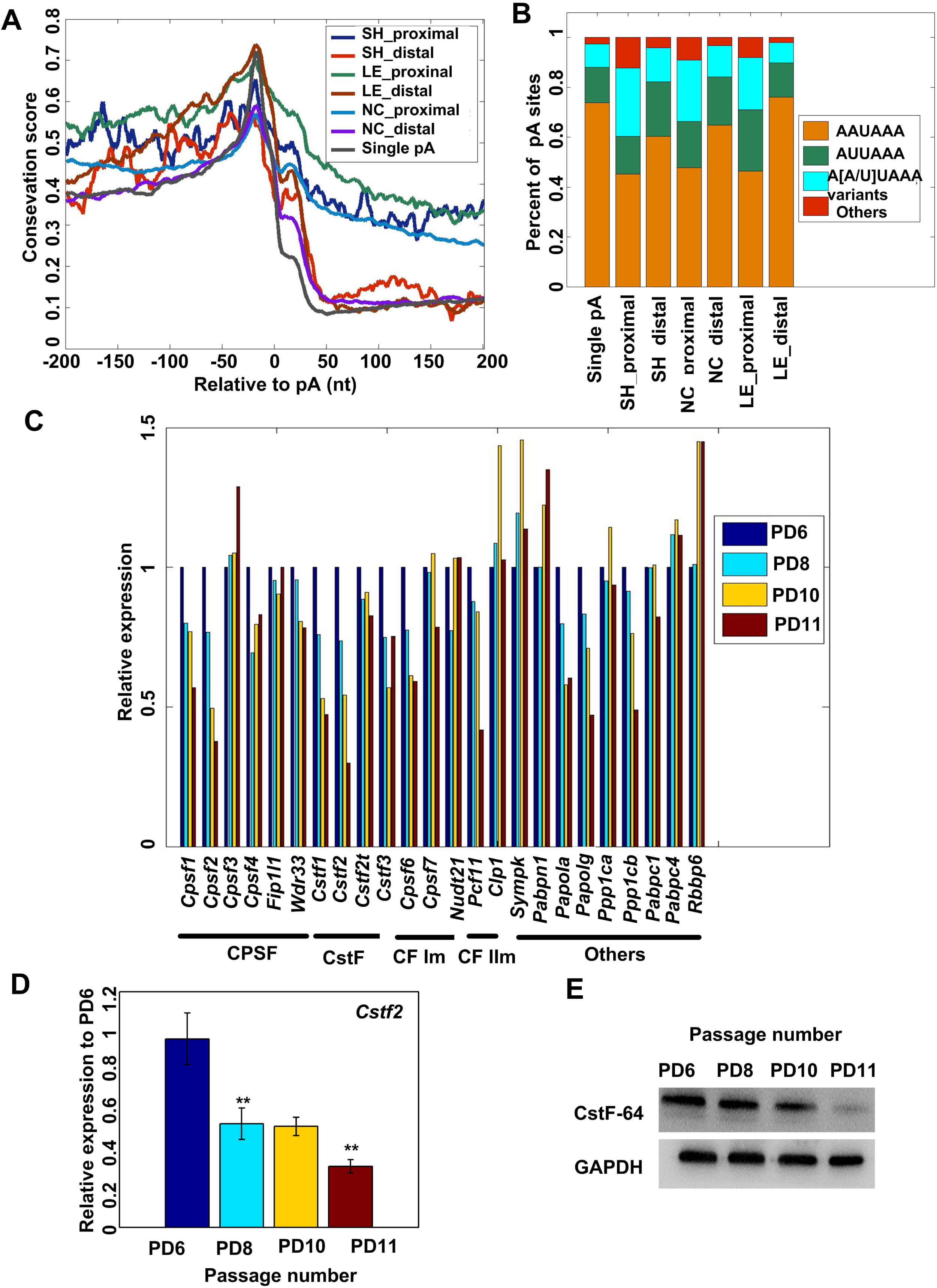
Potential mechanisms for APA regulation during replicative senescence of MEFs. (A) Comparison of conservation score in the 400 nt region surrounding different types of pAs. (B) Distribution of PAS sequences in the -40 to -1 nt region for different categories of pAs. (C) Gene expression of known cleavage and polyadenylation factors during replicative senescence of MEFs based on RNA-seq data. (D) Decreased expression level of *Cstf2* in PD6, PD8, PD10, and PD11 passages of MEFs were confirmed by qRT-PCR. (E) Detection of protein levels of CstF-64 (gene symbol is *Cstf2*) in PD6, PD8, PD10, and PD11 passages of MEFs by Western blot analysis. (**) *P* <0.01, two-tailed Student's t-test.

We next examined the possibility that RBPs control APA during cellular senescence. While the core factors in the 3' processing machinery play essential roles in the cleavage and polyadenylation (C/P) reaction, regulation of the level of expression has also been implicated in APA regulation (Zheng and Tian 2014). Therefore, the gene expression of 22 important poly(A) trans-factors were analyzed based on the MEF replicative senescence RNA-seq data (PD6, PD8, PD10, and PD11) (Xia et al. 2014). Consistent with the overall expression trend of RBPs, we observed global down-regulation of most poly(A) trans-acting factors during replicative senescence (Figure 8C), in which more genes tend to use a longer 3' UTR. This result is in agreement with a recently published study, in which global up-regulation of most poly(A) factors is associated with more 3'-UTR-shortening events in seven tumor types compared with normal tissues (Xia et al. 2014). Another survey of the constitutive components of the polyadenylation machinery and other candidates from the literature revealed several trans-acting factors that are significantly up-regulated in the cancer cell lines compared with non-transforming cells (Mayr and Bartel 2009).

Specifically, consistent with the notion that cellular senescence can be a defense system for cells on the way to becoming cancerous, we noticed that the mRNA level of Cstf2, which is the first core C/P factor that can regulate APA, was down-regulated during MEF cellular senescence (Figure 8C) and up-regulated in cancer cells (Takagaki et al. 1996; Mayr and Bartel 2009; Xia et al. 2014). Furthermore, Yao et al. showed knockdown of CstF64 (coded by *Cstf2* gene) led to more genes favoring distal pA sites, which contributed to lengthening of 3' UTR in human HeLa cells (Yao et al. 2012). We thus validated the mRNA expression and protein production of *Ctsf2* gene during MEF cell senescence. qRT-PCR (Figure 8D) and Western blot analysis (Figure 8E) confirmed that *Cstf2* showed a reduced trend of mRNA and protein expression, which may have contributed to 3' UTR lengthening during cellular senescence. Notably, Li et al. found that *Pcf11* could enhance usage of proximal pAs, and *Pabpn1* could promote usage of distal pAs by using siRNA knockdown coupled with deep sequencing (Li et al. 2015). Interestingly, the expression of *Pcf11* was down-regulated and *Pabpn1* was up-regulated during replicative senescence of MEFs, consistent with the global 3' UTR lengthening pattern. Taken together, we provided evidence that a change in expression of cleavage and polyadenylation-related factors, such as CstF64, Pcf11, and PABPN1, may regulate global 3' UTR lengthening during MEF cellular senescence, although the speculation deserves further experimental validation in MEF cells.

## Discussion

An increasing number of reports demonstrated the prevalence of global APA regulation in a wide range of biological processes and pathologic diseases (Ji et al. 2009; Mayr and Bartel 2009; Xia et al. 2014). Cellular senescence is regarded as a cancer prevention mechanism and a contributing factor for individual ageing (Campisi 2001; López-Otín et al. 2013). Increasing evidences support that cancer cells favor proximal pAs than normal cells (Mayr and Bartel 2009; Xia et al. 2014). Nevertheless, whether or not global APA regulation exists in senescent cells is completely unclear. We thus applied our PA-seq method to profile global APA events by using both MEF replicative senescence and VSMCs from rats with different ages as model systems (Busuttil et al. 2003; Parrinello et al. 2003; Di Micco et al. 2008; Manning and Kumar 2010). Interestingly, global 3' UTR lengthening during both senescent mouse and rat cells was observed, in contrast to global shortening of 3' UTR in tumor cells (Figure 3–4). Moreover, genes that prefer to use distal pAs in both senescent MFEs and VSMCs from elder rat show global lengthening of 3' UTR have a decreased trend of mRNA expression (Figure 4), consistent with the observation that genes prefer using proximal pAs in cancer cells have increased mRNA level ( Mayr and Bartel 2009; Xia et al. 2014). Furthermore, many genes were found to have opposite pAs usage preference between senescent and cancer cells (Supplemental Figure S10) (Xia et al. 2014). There are 82 and 166 genes preferred to use distal pAs in senescent MEFs and rat VSMCs, respectively, favored proximal pAs in seven tumor types (Supplemental Figure S10). Strikingly, 35 genes used longer 3' UTR in both senescent MEFs and rat VSMCs preferred shorter 3' UTR in multiple cancer cells (Supplemental Figure S10D). These findings support a model in which interaction between condition-specific *trans-acting* factors and dynamic changes of 3' UTR length determined by APA contributed to two opposite biological processes such as cellular senescence and tumor development.

Genes with longer 3' UTR in both senescent MEF and elder rat VSMCs are enriched in common pathways, including ubiquitin-mediated proteolysis (Vernace et al. 2007; Deschenes-Simard et al. 2014), regulation of the actin cytoskeleton (Gourlay and Ayscough 2005; Amberg et al. 2012), cell cycle (Chandler and Peters 2013), and the wnt signaling pathway (Hofmann et al. 2014; Lezzerini and Budovskaya 2014), which were confirmed to be involved in cellular senescence (Figure 5 and Supplemental Figure S9). Interestingly, the genes preferred to use proximal pAs and had increased levels of expression in cancer cells are also enriched in ubiquitin-mediated proteolysis and regulation of actin cytoskeleton pathways (Xia et al. 2014). These findings implied that APA regulation of genes involved in these two pathways may play a vital role in deciding cell fate, such as senescence or cancer.

Proteostasis is maintained by the proteostasis network that regulates protein synthesis, folding, trafficking, aggregation, disaggregation, and degradation (Powers et al. 2009). Of note, the identification of shared “ubiquitin-mediated proteolysis” pathway between senescent MEFs and VSMCs from elder rat is of interest as this machinery plays a key role in protein homeostasis, and functionality declines with age (Koga et al. 2011). In addition, dysfunction of the ubiquitin-proteasome pathway has been involved in pathogenesis of various neurodegenerative diseases, including Alzheimer's disease (AD) and Huntington disease (HD) (Bossy-Wetzel et al. 2004; Li and Li 2011). Specifically, NEDD4, an E3 ubiquitin ligase regulated by APA in senescent MEFs and VSMCs of elder rat (Figure 5C), was shown to delay cellular senescence by degrading PPAR*γ* and eventually up-regulating SIRT1 (Han et al. 2013). Interestingly, *Nedd4* was also shown to be a potential proto-oncogene that negatively regulated the phosphatase PTEN, a conserved component of the insulin/insulin-like growth factor-1-signaling (IIS) cascade, via ubiquitination (Wang et al. 2007). In addition, Cul1, which belongs to the Skp1-Cul1-F-Box E3 ligase complex, was also involved in regulating the lifespan of *Caenorhabditis elegans* (Ghazi et al. 2007). Furthermore, the Cul1 complex has been shown to inhibit the activity of FOXO proteins by catalyzing their degradation (Huang et al. 2005). *Cul1* can also promote cell-cycle progression at the G1-S phase transition through regulation of the abundance of cyclin E (Dealy et al. 1999). Interestingly, the longer 3' UTR of the *Cul1* gene, which was preferred to be used in senescent MEFs (Figure S11A-B), has a significantly lower stability than the shorter 3' UTR (*P*=3.5 × 10^−3^, *P*=1.5 × 10^−3^ and *P*=2.2 × 10^−3^ for 4, 8, 12 hours after actinomycin D treatment, respectively, two-tailed t-test; Supplemental FigureS11C). To examine how different 3' UTR isoforms of *Cul1* influence protein expression, the shorter, longer, and longer with mutated proximal PAS 3' UTRs (3' UTR_S, 3' UTR_L, and 3' UTR_M, respectively) were fused to a dual-luciferase reporter assay system, respectively. Both 3' UTR_L and 3' UTR_M yielded lower luciferase activity than the construct containing only the common 3' UTR region (3' UTR_S) in the NIH3T3 cells (*P*<0.04 and *P*<0.02, respectively, two-tailed t-test; Supplemental Figure S11D). The above data further support the idea that APA may be involved in cellular senescence via affecting the mRNA stability and protein translation efficiency of genes in senescence-related pathways.

Gourlay et al. demonstrated that elevated actin dynamic can increase the lifespan of yeast by more than 65% (Gourlay et al. 2004). In addition, CDK5 activation was up-regulated in senescent cells, which further accelerated actin polymerization and induced senescence and the senescent shape change by reducing GTPase Rac1 activity (Alexander et al. 2004). Florian et al. found that an unexpected shift from canonical to non-conical wnt signaling, caused by elevated expression of *Wnt5a* in aged haematopoietic stem cells (HSCs), leading to stem cell ageing in mice (Florian et al. 2013). Interestingly, we found both *Rac1* and *Wnt5a* tended to use distal pAs in senescent MEFs and VSMCs from elder rat. Collectively, our study revealed that APA regulation may be involved in several signaling pathways, which are likely implicated in cellular senescence.

Notably, there are two major categories of cellular senescence, including replicative senescence and stress-induced premature senescence (SIPS) (Munoz-Espin and Serrano 2014). Here we proved that MEFs replicative senescence underwent global 3' UTR lengthening. VSMCs derived from old rat are likely a combination of replicative senescence and varieties of stress-induced senescence. Although we discovered that more genes use a longer 3' UTR, whether or not SIPS itself will induce global 3' UTR lengthening remains elusive. Thus, more cellular senescent models are required to fully understand the prevalence and functional relevance of 3' UTR lengthening, which serves as an important post-transcriptional gene regulation mechanism. Importantly, PA-seq and RNA-seq can only manifest the steady-state level of mRNAs, and nascent transcript sequencing, such as GRO-seq (Core et al. 2008) and Bru-seq (Paulsen et al. 2014), may tell us whether or not the genes tended to use distal pAs during transcription in senescent cells. In addition, in this study we only predicted the potential binding sites for miRNAs and RBPs in 3' UTR regions. miRNA expression data and CLIP-seq (Yeo et al. 2009) or PAR-CLIP seq (Hafner et al. 2010) data for RBPs can help us further investigate how miRNAs and RBPs shape the mRNA fate after transcription. Furthermore, TAIL-seq, which allows us to measure poly(A) tail length at the genomic scale, is also a potent tool that can be used to investigate dynamic regulation of mRNA stability and translational control (Chang et al. 2014). Recently, Velten et al. developed BAT-seq to analyze single-cell isoform data, and found that single cells from the same state differed in the usage of APA isoforms (Velten et al. 2015). In this study, we only investigated the pA usage of genes from a cell population. Whether or not a more senescent cell is more prone to use distal pA can be further investigated using the same or a comparable method. In conclusion, the above results provide evidence to support the intrinsic link between APA and senescence in multiple species, although the functional validation and an underlying mechanism deserve further investigation.

## Methods

### Cell isolation and cultivation

Primary MEFs were isolated from embryo of a 12.5–14 day pregnant C57BL/6 mouse, according to the method described previously (Todaro and Green, 1963). Cells were cultured in DEME (GIBICO), supplemented with 10% fetal bovine serum (FBS; GIBCO) and 1% penicillin/streptomycin (GIBICO) at 37 °C in a humidified incubator with 5% CO_2_. To develop replicative senescence, confluent MEFs were evenly transferred into new dishes and the cells were cultured until confluent again to generate one PD. MEFs were maintained in a 100-mm dish, and in this study, confluent status of cells means almost confluent culture conditions without contact inhibition. The number (n) of PDs was calculated using the equation, where Ne and Ns are the number of cells at the end of cell culture and those seeded at the start of one passage, respectively. The number of PDs and total culturing period were monitored one time every 3 days until MEFs completely stopped proliferating. TRIzol reagent (Invitrogen) was used to extract total RNA from MEFs. Total RNA were also isolated from aortic vascular smooth muscle cells (VSMCs) in rats of different ages (2-week and 2-year) by TRIzol reagent.

### Cell cycle analysis by FACS

MEFs were grown in 100-mm dishes and harvested with 1 ml of 0.25% trypsin when 70–80% confluent. Then, the cells were washed twice with ice-cold 1× PBS, fixed with 70% ethanol for 2 hours (h) in 4 °C, and washed again with the same PBS before penetration with 1% Triton X-100 plus RNase A (40 U/ml) and stained with propidium iodide (50 μg/ml) on a shaking platform at 37 °C. When staining was completed, DNA content was measured on a FACS Calibur HTS (Becton Dickinson). The percentage of diploid cells in the G1, S, and G2/M phases were analyzed using Summit version 5.0 software (Dako).

### SA-β-gal staining

SA-β-gal activity, a classic biomarker of senescent cells, was also monitored at the same time for all growth stages of MEFs. The culture medium was removed from 12-well plates and the cells were washed once with 1 ml of 1× PBS and fixed with 0.5 ml of fixative solution for 10–15 min at room temperature. While the cells were in the fixative solution, the staining solution mix (staining solution, staining supplement, and 20 mg/ml of X-gal in DMSO) was prepared according to the manufacturer’s instructions (Biovision, cat. no. K320–250). After twice-washing with 1 ml of 1× PBS, cells were incubated with the aforementioned mix at 37 °C overnight or 12 h. Then, the cells were observed under an inverted microscope (Leica).

### qRT-PCR and Western blot

MEFs were harvested at 70%-80% confluence. Total RNA and protein were isolated with TRIzol reagent according to the provided manual (Invitrogen). For qRT-PCR analysis, RNA was reverse-transcribed into cDNA using random primers, and mRNA levels of *Mki67, p16*, and *Cstf2* were measured by qPCR (Roche LightCycler; see Supplemental Table S10 for primer information). For Western blot analysis, protein was resolved in 1% SDS, subjected to SDS-PAGE, transferred to nitrocellulose membranes, and incubated with primary antibodies of interest (p16, 1:500–1000, cat. no. sc-1207, Santa Cruz Biotechnology; Cstf2, 1:1000, cat. no. ab64942, Abcam; GAPDH, 1:2000–3000, cat. no. sc-32233, Santa Cruz Biotechnology) overnight at 4 °C. On the second day, the blots was washed three times in PBST (1× PBS + 0.1% Tween-20) and incubated with the diluted secondary antibodies (IRDye800CW goat anti-mouse, 1:10000, cat. no. 926–32210, LI-COR) for 1–2 h at room temperature. Then, the protein blots were washed three times with PBST before visualization under an Odyssey infrared imaging system (Odessey, LI-COR).

### PA-seq and RNA-seq library construction

The PA-seq libraries were constructed by following the protocol previously published by Ni et al. (Ni et al. 2013). The dUTP-based strand-specific RNA-seq libraries were constructed by following the protocol of Parkhomchuk et al. (Parkhomchuk et al. 2009). Both library types were sequenced by an Illumina HiSeq2000 platform in a paired-end 2×101 bp manner.

### Paired-end mapping and mRNA abundance estimation

Paired-end reads from PA-seq were subjected to strand correction according to ‘TTT’ at the beginning of the reads, as previously described (Ni et al. 2013). FastQC software was used for quality control of PA-seq and RNA-seq data (http://www.bioinformatics.bbsrc.ac.uk/projects/fastqc/). Processed raw data were then aligned to mouse genome (version mm9) or rat genome (version rn5) using Tophat2 (Kim D et al. 2013). Reads were mapped uniquely using the “-g 1” option. Bowtie was used to achieve a good approximation for -r and --mate-std-dev parameters, which were needed for Tophat2 (Langmead 2010). Combined counts of all pA clusters mapped to annotated PA, 3' UTR, and extended 3' UTR regions (number of Tags Per Million [TPM] reads) were used to evaluate the level of expression of these genes with APA regulation. RNA-seq reads were also aligned to the genome in a similar way and Cufflinks was used to assemble transcripts, estimate the mRNA abundance (FPKM) by following the instructions published previously (Trapnell et al. 2012).

### Peak calling

After mapping, the information of cleavage and polyadenylation sites (pAs) was extracted. All aligned reads were pooled together so that a unified peak-calling scheme could be applied. F-seq (Boyle et al. 2008) was used for peak calling of pAs by default parameter setting except that the feature length was set to 30. We resized the pA clusters to the shortest distance that contained 95% of the reads according to our previous publication (Ni et al. 2013). To filter internal priming, we removed pA clusters with 15 ‘A’ in the 20 nucleotides region downstream of peak mode or with continuous 6 ‘A’ in the downstream of peak mode. Peaks with fewer than 20 supporting reads were removed. RefSeq annotation was used for the genomic location analysis. We classified the genome into eight groups and mapped the pA peaks to these groups using the following hierarchy: PA > 3' UTR > extended 3' UTR > exon > intron > 5' UTR > TSS > promoter. Specifically, PA was defined as an annotated pA site plus upstream and downstream 10 bp. Extended_3' UTR denoted downstream 5 kb of 3' UTR by following the previous study (Ni et al. 2013). Note that the peaks mapped to an extended 3' UTR region should land in an intergenic region and not overlap with any known transcripts. Transcription start site (TSS) represented the exact transcriptional start site plus upstream and downstream 10 bp. The promoter was defined as the upstream 250 bp of TSS, which should not overlap with any known transcripts.

### Motif analysis

MEME motif enrichment analysis was used to identify the enriched motifs in the upstream (-40 nt) of our identified pAs with a motif length set to 6, which is equal to the length of PAS (Bailey et al. 2009). In addition, the frequencies of matches of 15 motifs identified by Hu et al. in different categories of pAs were defined by polya_svm (Hu et al. 2005; Cheng et al. 2006). Notably, only highly similar matches of motifs were considered (> 75th percentile of all possible positive scores) (Cheng et al. 2006).

### Comparison of APA profiles between different passage of MEF cells

The 3' UTR switching for each gene among the four passages was detected by measuring the linear trend similarly described as previously reported (Fu et al. 2011; Li et al. 2012). The genes with significant *P* value corresponding to a false discovery rate (FDR) cutoff of 5%, estimated by Benjamini-Hochberge method with R software, were considered as significantly different among different passages. In specificity, a FDR value less than or equal to 0.05 with positive TSI (tandem 3' UTR isoform switch index, same as the previous study (Li et al. 2012)) implies a lengthening 3' UTR across the different passages of cellular senescence; FDR value less than or equal to 0.05 with negative TSI implies a shortening 3' UTR.

### Pathway enrichment analysis

Database for Annotation, Visualization and Integrated Discovery (DAVID) (Huang et al. 2009) was used for the pathway enrichment analysis and the Kyoto Encyclopedia of Genes and Genomes (KEGG) database was selected.

### Prediction conserved binding sites of miRNA and RBP in 3' UTRs

The sequences of all mature miRNAs of mouse were downloaded from the miRBase database (release 21); 8mer, 7mer-m8, and 7mer-1A targets denote conventional miRNA seed matches (Hansen et al. 2013). Conserved target sites were identified using multiple genome alignments obtained from UCSC Genome Browser (mm9, 30-way alignment), requiring a perfectly aligned match in mice, rats, humans, and dogs, as in the paper of Sandberg et al. (Sandberg et al. 2008). Highly-conserved RBP binding sites were predicted by RBPmap online tool (Paz et al. 2014).

### Cluster analysis of RBPs

Clustering of genes expression level was done using the Cluster software (http://rana.lbl.gov/EisenSoftware.htm). K-means clustering approach based on euclidean distance was used to classify the various types of gene profiles. The cluster result was then visualized by Treeview software (http://taxonomy.zoology.gla.ac.uk/rod/treeview.html).

### Conservation analysis surrounding pAs

Conservation scores in the 400 nt surrounding different types of pAs were downloaded from UCSC genome browser (http://hgdownload.soe.ucsc.edu/goldenPath/mm9/phastCons30way/). The conservation score was measured by phastCons score based on 30 vertebrates (Thomas et al. 2003).

### Data Access

Both PA-seq and RNA-seq raw data can be found at the NCBI Sequence Read Archive (SRA) with submission number SRP065821.

## Acknowledgments

We are grateful to Dr. Haijian Wang for providing the psiCHECK-2 Vector, Professor Hongyan Wang for instrument support of luciferase assay and Professor Li Jin for insightful suggestions in bioinformatics analyses. This work was supported by National Key Basic Research Program of China (973 program: 2013CB530700 and 2015CB943000), National Science Foundation of China (31271348 and 31471192), Research and Innovation Project of Shanghai Municipal Education Commission (14ZZ007), and the 111 Project of China (B13016). We thank Genergy Biotech (Shanghai) Co., Ltd. for the deep sequencing service.

